# Insights on the assembly rules of a continent-wide multilayer network

**DOI:** 10.1101/452565

**Authors:** Marco A. R. Mello, Gabriel M. Felix, Rafael B. P. Pinheiro, Renata L. Muylaert, Cullen Geiselman, Sharlene E. Santana, Marco Tschapka, Nastaran Lotfi, Francisco A. Rodrigues, Richard D. Stevens

## Abstract

How are ecological systems assembled? Here, we aim to contribute to answering this question by harnessing the framework of a novel integrative hypothesis. We shed light on the assembly rules of a multilayer network formed by frugivory and nectarivory interactions between bats and plants in the Neotropics. Our results suggest that, at a large scale, phylogenetic trade-offs separate species into different layers and modules. At an intermediate scale, the modules are also shaped by geographic trade-offs. And at a small scale, the network shifts to a nested structure within its modules, probably as a consequence of resource breadth processes. Finally, once the topology of the network is shaped, morphological traits related to consuming fruits or nectar determine which species are central or peripheral. Our results help understand how different processes contribute to the assemblage of ecological systems at different scales, resulting in a compound topology.

## Introduction

Since Darwin’s “tangled bank” metaphor^1^, one of the most important quests in ecology has been to unveil the assembly rules of ecological systems^2^. Different study models have been used in an attempt to generate unifying principles, from sets of species (i.e., communities^3^) to systems formed by species interactions (i.e., networks^4^). Knowing those rules is crucial for understanding the architecture of biodiversity^5^, restoring degraded environments^6^, and controlling emerging diseases^7^, among other applications. However, identifying those rules remains one of the main unsolved challenges in ecology^8^.

Major advances in network science have shed light on some assembly rules that govern interaction systems^9–11^. These breakthroughs permitted the ecological and evolutionary analysis of monolayer networks formed by a single interaction type. Since then, there has been much debate concerning the prevalent topology among interaction networks (nested or modular) and which processes should generate those patterns (niche or neutral). Early evidence suggested that antagonistic networks should be predominantly modular, while mutualistic networks should be nested^12^. However, recent studies suggest that those topological archetypes are not exclusive to particular interaction types^13^, may occur in combination^14^, and depend on geographic and phylogenetic scales^15^.

A novel conceptual framework, termed “the integrative hypothesis of specialization” (IHS^16^), proposes that a balance between trade-offs^17^ at larger scales and resource breadth processes^18^ at smaller scales shapes host-parasite networks. The IHS, in its updated form^19^, is based on premises that can be extrapolated from parasites to consumers in general: (i) types of resources differ in their ability to be exploited by consumers; (ii) resources are more different from one another at larger than smaller scales; and (iii) an adaptation to exploit a resource helps exploit similar resources but becomes a maladaptation to exploit dissimilar resources.

Using the framework from the IHS and new models of multilayer networks^20^, here we aimed to understand the assembly rules of a system formed by bats and plants that interact with one another through frugivory and nectarivory in the entire Neotropical region. From the IHS, we deduced that different processes should shape the bat-plant network at different scales. If this is true, firstly, there should be strong phylogenetic and geographic trade-offs in the network studied, as it contains two interaction types and high phylogenetic diversity (one large bat family and several plant families^21^), distributed over an entire biogeographic region. These trade-offs should lead to strongly separated layers (large scale) and modules (intermediate scale). However, within the modules (small scale) resource breadth processes should lead to a nested structure, resulting in a compound topology: a modular network with modules internally nested. Secondly, considering that some bat species are able to feed both on fruits and nectar^22^, different organismal traits related to those diets^22,23^ should thus determine the relative importance of different bat species for the structure of each layer and for bridging layers.

Our results support the IHS as a good model to explain the topology of interaction networks. They also provide the first evidence of a compound topology in multilayer networks, with different processes operating at different scales.

## Results

The Neotropical bat-plant multilayer network analyzed here (Fig. 1a) is hyper-diverse and massive. It is composed of 439 plant species, 73 bat species, 911 links of frugivory, 301 links of nectarivory, and 18 dual links (i.e., links of both frugivory and nectarivory between the same bat and plant species). The frugivory layer contains 307 plant species and 56 bat species, while the nectarivory layer contains 139 plant species and 39 bat species. The 18 dual links were made between 10 bat species and 8 plant species.

**Figure 1.**
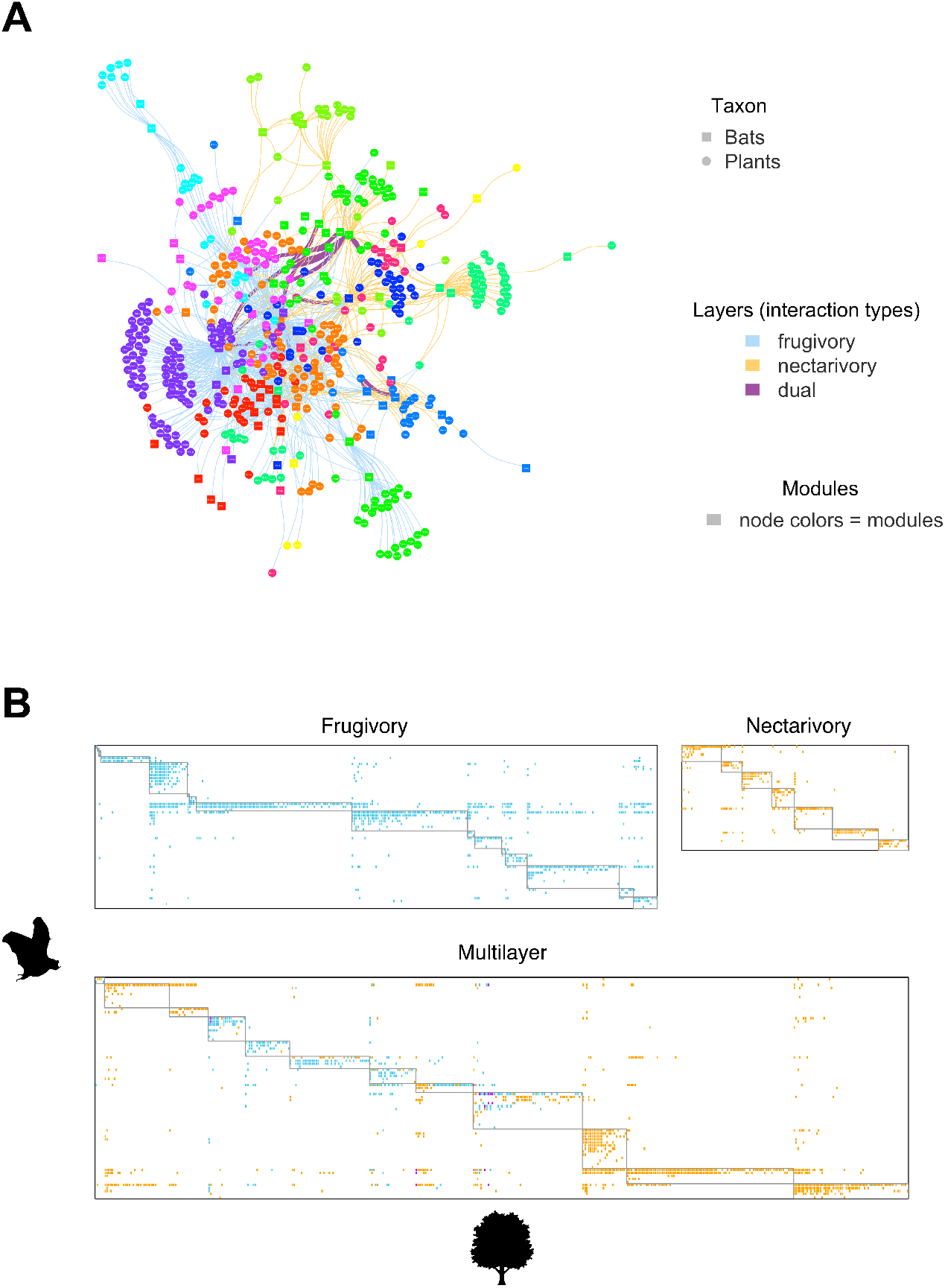
The multilayer bat-plant network, built for the entire Neotropical Region based on interactions of frugivory and nectarivory recorded in the wild, showed a strong separation between interaction types (layers) and guilds (modules). **A**. Multilayer graph; the layers represent interactions of frugivory (blue), nectarivory (orange), and dual interactions (purple, i.e., interactions of both types between the same bat and plant species). Bat species are represented as squares, plant species as circles, and interactions as lines. Node colors represent modules detected in the network using the DIRT_LPAwb+ algorithm. **B**. Multilayer matrix; bat species are represented in the rows, plant species in the columns, and filled cells represent interactions (same colors as in the graph); boxes represent the modules found. See the full-sized graph with species labels in Supplementary Data 1.

As predicted, the studied network showed a compound topology (Table 1, Fig. 1b). The modularity score for the whole multilayer structure (M = 0.53, Z_free_ = 49.18, P <0.001) was much higher than expected by the free null model, which does not consider the network’s modular structure (see Methods for explanations of the null models). The same was observed for the frugivory (M = 0.48, Z_free_ = 44.44, P <0.001) and nectarivory layers, respectively, using the free null model (M = 0.63, Z_free_ = 24.94, P <0.001). In contrast, the entire multilayer structure was slightly nested (NODF = 0.18, Z_free_ = 4.72, P_free_ < 0.001), as well as the frugivory (NODF = 0.29, Z_free_ = 7.12, P_free_ < 0.001) and nectarivory layers (NODF = 0.16, Z_free_ = 2.39, P_free_ < 0.013). In other words, the studied multilayer network is both modular and nested at the same time, but the modular structure is stronger than the nested structure at larger scales.

**Table 1.** The multilayer network presented a compound topology, with a modular structure that comprises internally nested modules. Scores of modularity (M) and nestedness (NODF) for the entire multilayer matrix and its layers, including NODF scores calculated between species of the same module (sm) and of different modules (dm). The scores were calculated for the studied matrix (Obs), and also randomized according to the free and restricted null models. P-values (P) were estimated based on a Monte Carlo procedures run for each null model (1,000 iterations), which lead to expected scores (E) and Z-scores (Z). The free null model randomizes the entire matrix, whereas the restricted null model conserves its modular structure. We did not run the fixed null model for modularity. All scores were standardized varying from 0 to 1. Significance level (α): 0.05.

|  | **Obs** | **E<sub>free</sub>** | **Z<sub>free</sub>** | **P<sub>free</sub>** | **E<sub>rest</sub>** | **Z<sub>rest</sub>** | **P<sub>rest</sub>** |
| --- | --- | --- | --- | --- | --- | --- | --- |
| **Frugivory layer** |  |  |  |  |  |  |  |
| Mod | 0.48 | 0.35 | 44.45 | 0.001 | NA | NA | NA |
| NODF | 0.29 | 0.22 | 7.00 | 0.001 | 0.23 | 6.44 | 0.001 |
| NODF<sub>sm</sub> | 0.60 | 0.19 | 34.69 | 0.001 | 0.43 | 8.37 | 0.001 |
| NODF<sub>dm</sub> | 0.23 | 0.22 | 0.92 | 0.179 | 0.19 | 4.04 | 0.002 |
| **Nectarivory layer** |  |  |  |  |  |  |  |
| Mod | 0.63 | 0.47 | 24.95 | 0.001 | NA | NA | NA |
| NODF | 0.16 | 0.13 | 2.39 | 0.013 | 0.13 | 3.02 | 0.003 |
| NODF<sub>sm</sub> | 0.55 | 0.13 | 37.41 | 0.001 | 0.35 | 8.41 | 0.001 |
| NODF<sub>dm</sub> | 0.09 | 0.13 | -4.82 | 0.999 | 0.09 | -0.56 | 0.710 |
| **Multilayer** |  |  |  |  |  |  |  |
| Mod | 0.53 | 0.38 | 49.18 | 0.001 | NA | NA | NA |
| NODF | 0.18 | 0.15 | 4.73 | 0.001 | 0.15 | 6.14 | 0.001 |
| NODF<sub>sm</sub> | 0.55 | 0.14 | 53.32 | 0.001 | 0.40 | 8.85 | 0.001 |
| NODF<sub>dm</sub> | 0.13 | 0.15 | -2.23 | 0.994 | 0.11 | 3.58 | 0.001 |

Corroborating this result, nestedness between species of different modules (NODF_DM_) was lower than expected by the free null model in the nectarivory layer and the multilayer network but, interestingly, equal to the expected value in the frugivory layer. This result suggests that the modules impose greater constraints to nectarivory than to frugivory interactions. Furthermore, nestedness in general (NODF), between species of the same module (NODF_SM_), and between species of different modules (NODF_DM_) was higher than expected considering the modular structure of the multilayer network and its layers. The exception was the nectarivory layer, in which species of different modules (NODF_DM_) show higher nestedness than expected given the modules.

Geographic co-occurrence and phylogeny of bat species were also important predictors of the network’s compound structure. Most bat species analyzed have small geographic ranges, while a few are broadly distributed. The species with the smallest range was *Lonchophylla bokermanni* (23,309 km^2^), whereas the species with the largest range was *Sturnira lilium* (17,327,789 km^2^). Mantel tests found no correlation between the geographic co-occurrence and phylogenetic distances of bat species, which means that these bat clades are distributed in the Neotropical Region independently of their evolutionary origin (Figure 2a). Though we found a strong phylogenetic signal in the modules of the network (intermediate scale) we did not find such signal in the interactions within the modules (small scale). Nevertheless, the contrary was true for the geographic signal: it is strong at the scale of within-module interactions, but very weak in the modules (Figure 2a). We found these same general trends (Figure 2b-c) when we used partial Mantel tests to discount for mutual effects between the structuring factors and the scales.

**Figure 2.**
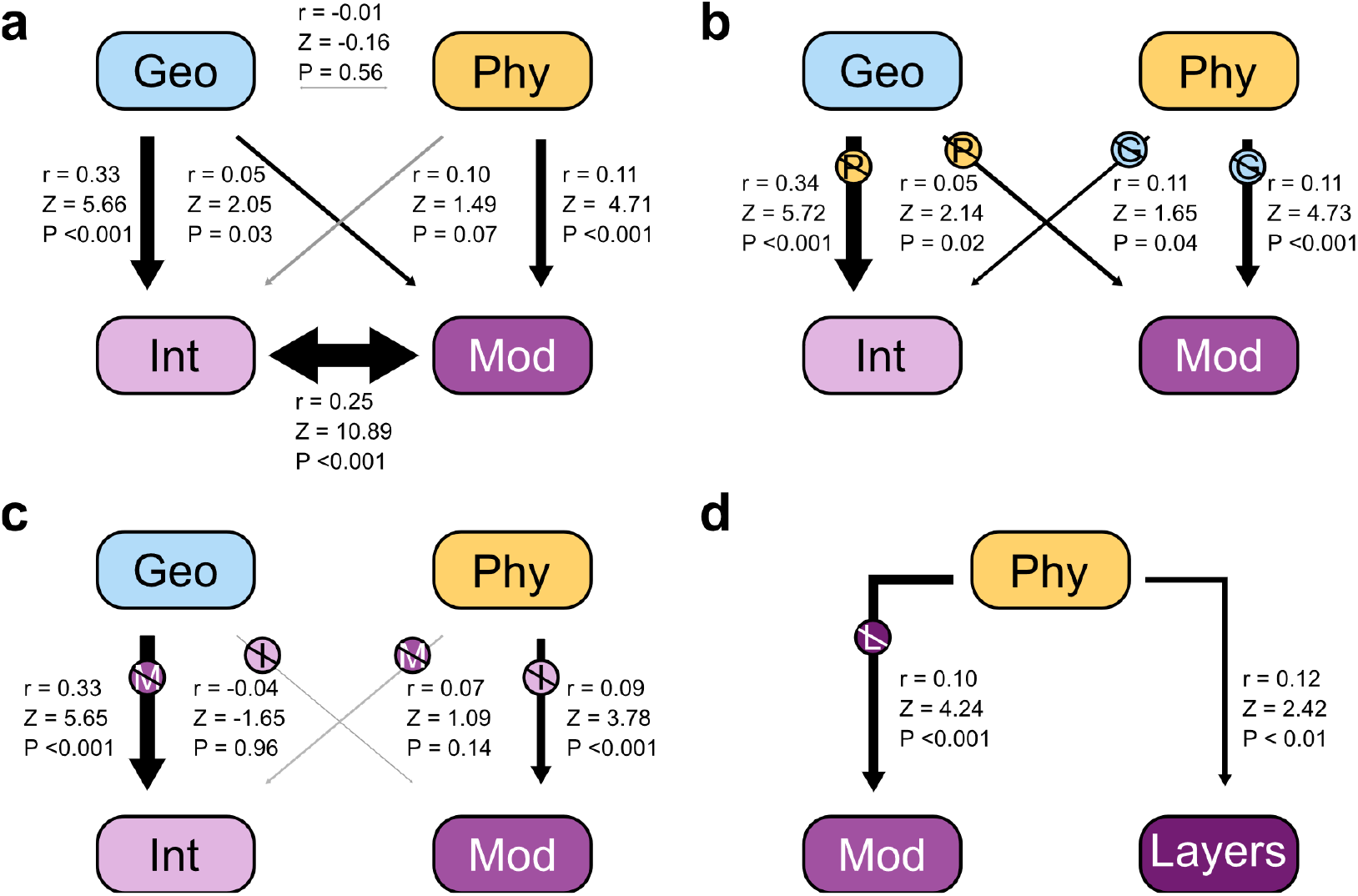
Phylogenetic (Phy) and geographic (Geo) signals at different scales of the multilayer network: interactions (Int) (small scale), modules (Mod) (intermediate scale), and layers (large scale). **A**. Results of Mantel tests for all the correlations between bat distances in phylogeny, geographic co-occurrence, interactions, and modules. **B**. We used partial Mantel tests to discount the mutual effects between phylogeny and geographic co-occurrence; therefore, when testing geographic signals at each scale, we conditioned the correlation on the phylogenetic distance, and vice-versa. **C**. We used partial Mantel tests but conditioning the correlations with distances in one scale on the distances in the other scale. **D**. We used a Mantel test to assess a phylogenetic signal in the layers of the network and then used a partial Mantel test to test the phylogenetic signal in the modules accounting for the distance between layers. Arrows in black represent significant correlations and in gray, non-significant correlations. Arrow width scaled by Z-scores. In partial Mantel tests, the crossed circle with a letter inside indicates on which distances the correlation tested (arrow) was conditioned (geographic – G, phylogenetic – P, modules – M, or interactions – I).

There was dependence between modules and interaction types (χ^2^ = 554.33, N = 12, P < 0.001), which means that some modules are formed mainly by nectarivorous bats and others by frugivorous bats. Additionally, we detected a phylogenetic signal in layer composition (Figure 2d), where some bat clades are preferentially nectarivorous while others are preferentially frugivorous, which corroborates the structuring power of phylogeny at a large scale. Then, because of the dependence between layers and modules, we tested and confirmed that the phylogenetic signal in the modules remains even when discounting the correlation with the layers (Figure 2d). We conclude that phylogeny structures the layers of the network (large scale) and the modules inside each layer (intermediate scale), and geographic co-occurrence structures the interactions within each module (small scale). Finally, there was no phylogenetic signal in bridge species, which make both interactions of frugivory and nectarivory (r = 0.04, Z = 0.75, P = 0.21).

Few centrality metrics presented significant correlations with one another, whereas most were only weakly correlated or not correlated (Supplementary Results 1). Centrality varied largely among all species. It varied also between layers in the case of bridge species (Figure 3). For these bridge species, there was no relationship between degree, betweenness centrality, closeness, or eigenvector centrality across layers (all P > 0.05, Table 2, Supplementary Results 2). However, bat species with larger degree, betweenness centrality, and eigenvector centrality in the frugivory layer had higher probabilities of being bridge species (all P < 0.05, Table 2, Supplementary Results 2). In the nectarivory layer, none of the centrality metrics explained the probability of a species being a bridge between layers.

**Figure 3.**
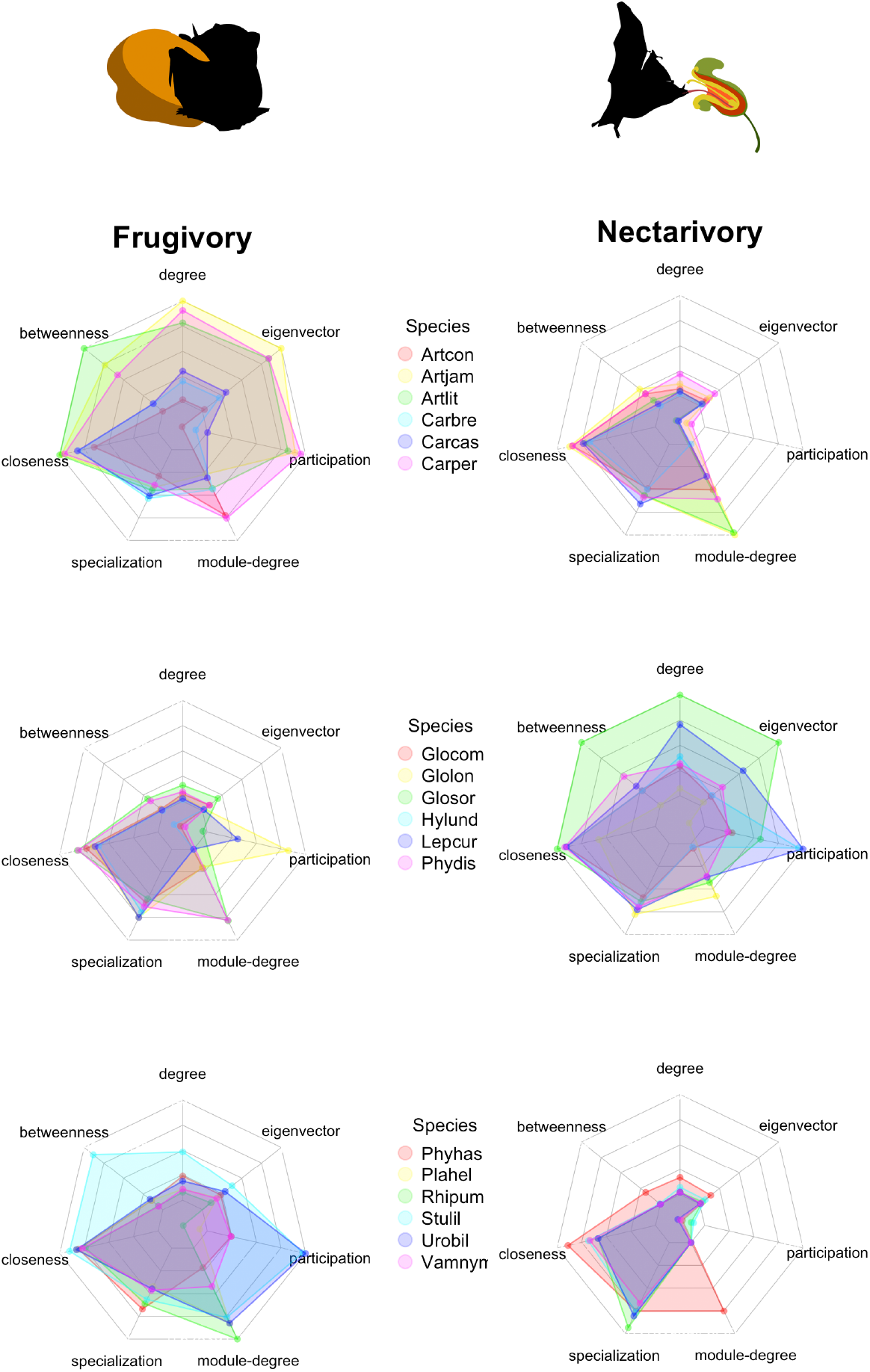
The centrality metrics varied largely in the same species between layers of the network (frugivory and nectarivory). Each axis of each spider chart represents a centrality metric calculated, and its original range of variation. Different bat species are represented by different colors. Only the most central species that occurred in both layers are presented here. Species codes were made using the first three letters of the genus and the first three letters of the epithet (e.g., Carper = *Carollia perspicillata*). See binomial nomenclature in Supplementary Data 1.

**Table 2.** The centrality of a bat species on one layer of the network did not predict its centrality on the other layer. However, the higher the centrality of a bat species in the frugivory layer, the higher its probability of being a bridge species (i.e., making interactions on both the frugivory and the nectarivory layers). Relationships between centrality metrics calculated in different layers of the network using GLMs. Significant P-values are highlighted in boldface. Significance of the models of the set 1 was estimated using F tests, while for the sets 2 and 3 we used χ² tests.

| Model | df | deviance | F | P |
| --- | --- | --- | --- | --- |
| 1. Centralities vs. layers |  |  |  |  |
| ndeg.frug ~ ndeg.nect | 20 | 0.027 | 0.269 | 0.610 |
| bet.frug ~ bet.nect | 20 | 0.003 | 0.033 | 0.857 |
| clo.frug ~ clo.nect | 20 | 0.002 | 2.208 | 0.153 |
| eig.frug ~ eig.nect | 20 | 0.018 | 4.870 | 0.832 |
| 2. Bridge species vs. frugivory |  |  |  |  |
| ndeg ~ bridge | 54 | 12.607 |  | **0.000** |
| bet ~ bridge | 54 | 16.125 |  | **0.000** |
| clo ~ bridge | 54 | 1.119 |  | 0.290 |
| eig ~ bridge | 54 | 14.940 |  | **0.000** |
| 3. Bridge species vs. nectarivory |  |  |  |  |
| ndeg ~ bridge | 41 | 0.073 |  | 0.787 |
| bet ~ bridge | 41 | 0.858 |  | 0.354 |
| clo ~ bridge | 41 | 1.759 |  | 0.185 |
| eig ~ bridge | 41 | 0.002 |  | 0.963 |

Legend: ndeg = normalized degree, bet = betweenness, clo = closeness, eig = eigenvector, frug = frugivory layer, nect = nectarivory layer. Significance level ( $\alpha$ ): 0.05.

Geographic range size did not affect the centrality of bat species. Among the morphological attributes, body size and bite force were the most important predictors of species’ centrality. For the frugivory layer, the latent variable analysis (N = 16, df = 29) indicated that eigenvector decreased with body size (coefficient = -0.524, P = 0.003), increased with bite force (coefficient = 1.585, P < 0.001), and was not explained by the other latent and indicator variables (Fig. 4a). For the nectarivory layer (N = 15, df = 29), eigenvector increased with body size (coefficient = 1.268, P < 0.001), decreased with bite force (coefficient = -1.841, P < 0.001), and was not explained by the other variables (Fig. 4b). For dual interactions, the model could not be calculated due to the small number of observations. Finally, considering the entire multilayer structure and a multilayer version of centrality (N = 18, df = 29), eigenvector increased with bite force (coefficient = 0.517, P = 0.013), and was not explained by the other variables (Fig. 4c).

**Figure 4.**
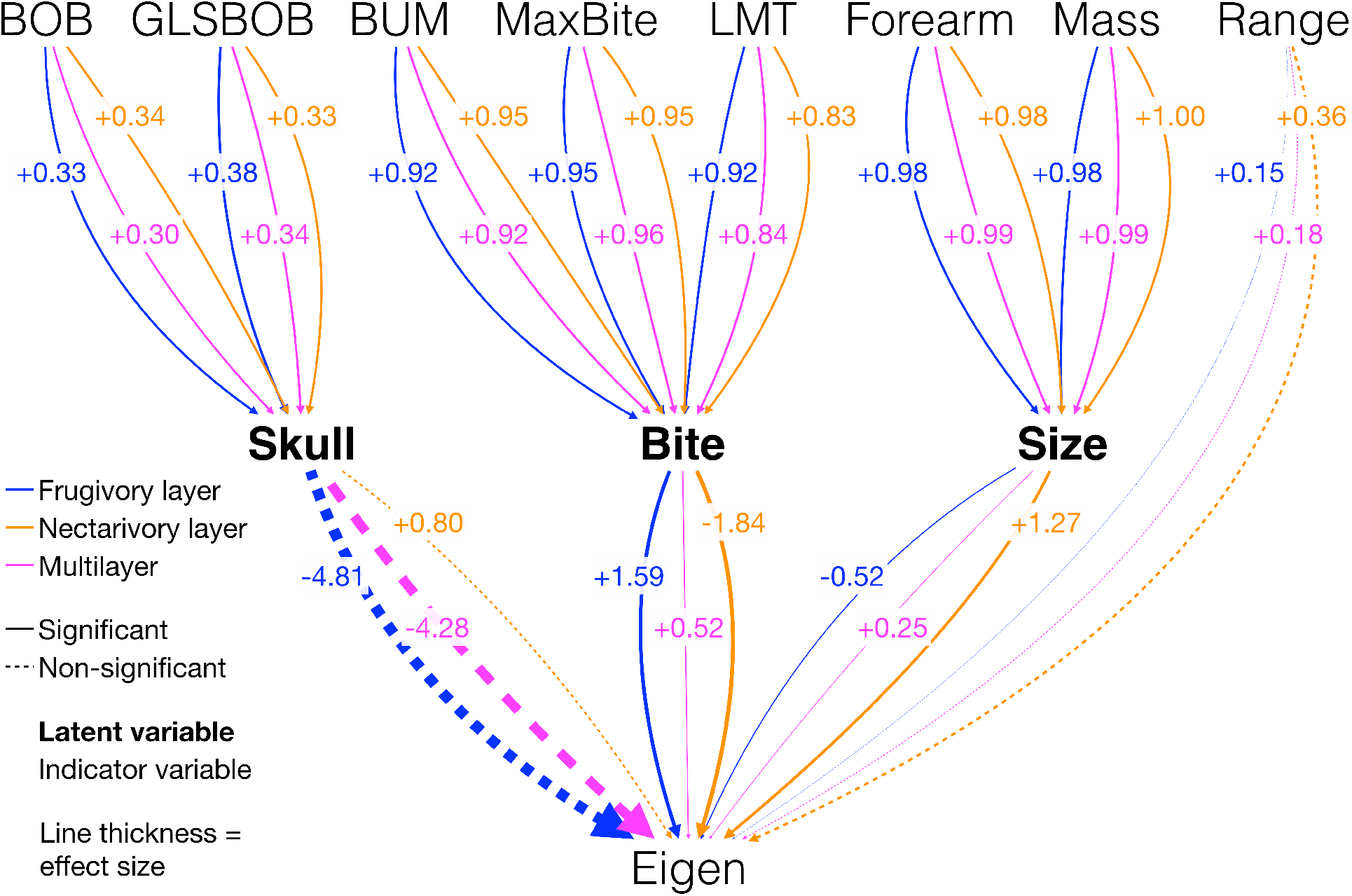
The eigenvector centrality (Eigen) of a bat species was determined by a combination of biological traits (indicators) related to morphology (the latent variables: Skull, Bite, Size) and geographic range size (Range). A bat species was more central in the frugivory layer, when it had a strong bite force (Bite) and small body (Size). In the nectarivory layer, larger bats (Size) with weak bite force (Bite) were the most central. In the complete multilayer structure, only bite force (Bite) was positively related to centrality. Numbers on the lines are the standardized coefficients of each path, and line thickness was drawn proportionally to this coefficient only for the latent variables (Skull, Bite, Size) and single indicator variable (Range). Significance was estimated only for those main variables. See full indicator names in Supplementary Results 1

## Discussion

Our analysis of a continent-wide multilayer interaction network shows that, in order to build a complex ecological system, a combination of processes operating at different scales is needed. This finding supports the integrative hypothesis of specialization (IHS^16,19^), which we here extend from parasite-host to plant-animal interactions.

Firstly, at large and intermediate scales, phylogenetic and geographic trade-offs generate a multilayered and modular structure. After the influence of those trade-offs, at a small scale, the modules of the network are internally nested and shaped first by geographic trade-offs. For sympatric species, this nested structure is probably a result of resource breadth processes^18^, neutral processes related to differences in abundance^24^, or universal processes observed in different kinds of complex networks such as preferential attachment^25^. Scale-dependence has been pointed out as a critical issue in biodiversity research^26^ and here we show that the same is true for species interactions. Secondly, after the network is shaped, biological traits determine how important each species is for the structure of each layer of the network. Those traits determine also which species bridge the layers by being frugivorous and nectarivorous at the same time.

Organismal attributes, such as body size and bite force, predict eigenvector centrality in a manner that is consistent with predictions from ecomorphological theory; species with greater performance are expected to have access to a broader array of ecological resources^27^. Bite force is a whole-organism performance trait that is tightly linked with the physical demands imposed by diet^28^. Specialized neotropical frugivores have evolved foreshortened rostra and large jaw adductors, which allows these species to have exceptionally forceful bites for their size and consume fruit across a broader hardness spectrum than species that have weaker bite forces^23,29,30^. Conversely, an elevated bite force is not a feeding performance requirement for nectarivores, to whom an elongated skull and thus weaker bite forces, and a larger body size may be an advantageous trait for accessing a broader array of flower sizes and types^31^.

Our results suggest that the dilemma of identifying the predominant topology among interaction networks (nested or modular) creates a false dichotomy. This interpretation is supported not only by our results, but by evidence from other recent studies^14,15,32^, which highlight that modularity and nestedness are states along a continuum^3^. Ecologists foresaw this continuum for interaction networks in the past^14^, and it seems applicable to other types of ecological systems, such as communities and metacommunities^3,33^. The IHS provides us with a mechanistic model that predicts this compound topology^16,19^. In addition, the evidence provided here also corroborates the importance of biological traits to the hierarchy of centrality in interaction networks^34,35^.

In conclusion, we found evidence that to build a continent-wide, hyper-diverse interaction network, we need different processes operating at different scales. Our findings integrate different debates in the ecological and parasitological literatures, and may also help understand the emergence of hierarchical structures in other complex systems, such as social and economic networks^36^.

## Methods

### Data set

The data set used in the present study came from the Bat-Plant Interaction Database^37^, which was partially published in a book on seed dispersal by bats^38^, and was later updated and used in other studies on ecological networks^34^. In the present study, we added new data on bat-flower interactions collected by the authors in Mexico, Costa Rica, and French Guiana, which were published in different papers. The list of data sources is presented in Supplementary Table 1.

### Network building

The original studies from which we sourced the bat-plant interaction data used a myriad of methods, ranging from mist-netting to roost inspection and direct observation. In addition, these studies varied in their focus, from single bat species or plant families, to whole bat-plant ensembles at a local scale^39^. Therefore, we decided to use binary data (i.e., presence or absence of interactions) to build the multilayer network, as it would be very complicated to integrate and standardize frequency data from different methods collected at different taxonomic scales. Furthermore, binary data are more adequate to assess fundamental ecological niches^40,41^, which is the case of our study. The multilayer network was compiled at the scale of the whole Neotropical Region. Henceforth, its nodes represent interactions across the entire geographic range of species of bats and plants, and not just single local populations. Its binary links (edges) thus represent dimensions of the fundamental niches of those species, and not their local realized niches.

On each layer of the network a bat species and a plant species were connected to each other by a link, whether an interaction of frugivory or nectarivory between them had been recorded in the wild. Several species make links of both types, and thus belong to both layers of the network. We call these “bridge species”. Furthermore, a few bat and plant species were connected to one another in both layers, making what we call here “dual links”. In other words, some bat species are both seed dispersers and pollinators of the same plant species. Consequently, the multilayer network contained two types of links: frugivory and nectarivory. Those link types were modeled as interconnected layers in the format of an edge list (Supplementary Data 1). See also Supplementary Methods 1, where we explain how the multilayer structure was modeled. Full Latin names of bats and plants are presented in Supplementary Data 1. Network science terms used here are explained in detail in Supplementary Table 2.

### Compound topology

#### Compound topology analysis

To test whether each layer and the aggregated network were formed by internally nested modules (compound topology, sensu^14^), we used a recently proposed protocol^19^, which is based on the following steps.

Step 1, find the best partition of a network and compare its modularity score to that expected by a given null model of interest^42^. Step 2, compute the nestedness (NODF) of the entire network and disentangle it into two components: nestedness between pairs of species of the same module (NODF_SM_) and nestedness between pairs of species of different modules (NODF_DM_). Step 3, compare the observed values of NODF_SM_ and NODF_DM_ to their values expected both in the absence (free null model) and in the presence (restricted null model) of the modular structure.

In a modular network, NODF_SM_ should be higher than expected when interactions are reshuffled regardless of the modular structure, *i.e*., following the free null model. The reason is that connectance of areas within the modules of the null matrices will be smaller than that of the real matrix, and NODF increases monotonically with connectance^43^. Therefore, to test whether interactions are more nested than expected given the modular structure, we compared the observed NODF_SM_ and NODF_DM_ to the values expected by a null model that conserves the modular structure (*i.e*., keeps the observed connectance values within and between modules in the null matrices).

#### Null models

On the one hand, the free null model produces null matrices of the same size, connectance, and species relative degrees. On the other hand, besides size, connectance, and relative degrees, the restricted null model also conserves the modular structure of the original matrix when generating the null matrices. This is made by weighting the *a priori* probability of interaction among species C_i_ and resource R_j_ (P_ij_) by the connectance of the matrix sub-area to which the cell M_ij_ belongs^19^.

For each layer, and the aggregated network, we generated 1,000 random matrices using the free null model and another 1,000 matrices using the restricted null model. Next, for each random matrix, we computed its overall NODF and decomposed it into NODF_SM_ and NODF_DM_ using the observed partitions of their corresponding real network. Finally, a Z-score was calculated as Z = [Value_obs_ – mean(Value_sim_)] / σ(Value_sim_), where Value_obs_ is the observed value of the metric and Value_sim_ represents the values of the metric in the randomized matrices. Observed and expected modularity values were also compared using Z-scores, but only for the free null model, as it does not make sense to compare observed and expected modularities with a null model that fixes the modules.

### Geographic and phylogenetic signals

We used a combination of analyses to detect the signals of the geographic distribution and of the phylogeny of bats at different scales of the multilayer network. In this analysis, we used only the bat species that belong to the main component of the network, whose distribution data were available in the IUCN red list global assessment (65 bat species). First, we computed five pairwise distance matrices for bat species: phylogenetic, geographic, interactions, modules, and layers.

To generate the phylogenetic distance matrix, we used the branch lengths in the most up-to-date, species-level phylogeny of phyllostomids^44^ (for 8 bat species not presented in the phylogeny we used an alternative approach, see Supplementary Methods 2). For the pairwise geographic distances, we used a measure of the overlap in the distribution of bat species recovered from IUCN databases. Interaction, module, and layer pairwise distances were calculated based on Jaccard Index (for details, see Supplementary Methods 2).

To test the signals, we performed a combination of Mantel and partial Mantel tests, and used the Z-Score as a measure of effect size (observed correlation minus the average correlation in randomized matrices, divided by the standard deviation). We tested the dependence between modules and layers of the network using a chi-squared test of independence. Lastly, we used a Mantel test to test for a phylogenetic signal in bridge species.

### Centrality and biological traits

We assessed the relative importance of each bat species to the structure of each layer or the entire multilayer network through the centrality metrics degree, closeness centrality, betweenness centrality, complementary specialization, within-module degree, participation coefficient, and eigenvector centrality. For details on their definition and calculation, see Supplementary Methods 1.

Using generalized linear models (GLMs) based on a quasi-Poisson distribution of errors, we tested whether the centralities of bat species in the frugivory and the nectarivory layers were correlated with one another. All models were checked for over- and sub-dispersion, and then tested with an analysis of variance (ANOVA).

To test for a correlation between centrality indices of bat species in each layer (frugivory and nectarivory) and the probability of a bat being a bridge species between the layers, we also used generalized linear models (GLMs). Since the response variable was binary (bridge species: yes or no), we used a binomial distribution of errors in those GLMs. We checked all models for overdispersion, and then tested them with a chi-squared test. These two first sets of statistical tests were conducted in R, using the package *lme4*^45^ (see Supplementary Results 2).

To test the relationship between body size, skull morphology, feeding performance, geographic range size, and centrality, we used a dataset on morphometric and performance traits of phyllostomid bats for the whole Neotropics, compiled by R. Stevens and S. Santana from published studies^23,29,46^. This dataset spans a large variety of morphometric and feeding performance traits, which were collected from wild animals and museum specimens using standardized methods^29^. As many of these are strongly correlated with one another, we relied on previous studies to select traits considered most relevant to feeding function in the context of frugivory and nectarivory (see Supplementary Results 1).

In relation to organismal traits, species with larger geographic range size are expected to have broader diets within their trophic niches (e.g., frugivory or nectarivory), as they cannot rely on specialized diets all over their distribuition^47,48^. Animals with larger body size are expected to have broader diets, as they have higher energy requirements than small-bodied animals^34,49^. Frugivorous bats are expected to bite more forcefully than nectarivorous bats, considering differences in hardness between solid and liquid diets^29^. Skull morphology is another important trait related to diet in bats, as frugivorous species tend to have shorter and broader skulls than nectarivorous species^22^.

As there should be complex direct and indirect paths of influence among body size, dietary morphology and performance, geographic range size, and centrality, we used a latent variable analysis (LaVaAn) to disentangle these relationships. In all models, the response variable, eigenvector centrality (eg.), was determined by three latent variables: body size (Siz), bite force (Bit), and skull morphology (Skl), and one single indicator: geographic range size (rng). The latent variable body size was composed of the exogenous variables body mass (Mss) and forearm length (Frr). The latent variable Bit was composed of the exogenous variables length of maxillary toothrow (LMT), breadth across upper molars (BUM), and maximum bite force (MBF). The latent variable Skl was composed of the exogenous variables breadth of braincase (BOB) and greatest length of skull (GLS). We built four similar models: one for the frugivory layer, one for the nectarivory layer, one for dual interactions (i.e., the same bat and plant species connected to one another in both layers), and one for the entire multilayer.

As not all bat species participate in both layers of the network, the sample size (N) of each test was smaller than the number of bat species analyzed in the present study. All statistical tests related to this prediction were carried out in R, using the package *lavaan*^50^ (significance level α = 0.05 for all tests).

## Acknowledgments

We are deeply grateful to all colleagues who carried out fieldwork in the Neotropics over several decades and collected the information used to build our data set. Paulo Guimarães Jr. and Thomas Lewinsohn discussed with us the assembly rules of ecological systems. Pedro Jordano, Carsten Dormann, and Katherine Ognyanova gave us invaluable tips on how to draw networks in R. Mark White and the StackOverflow community helped us build the model used in the latent variable analysis. MARM was funded by the Research Dean of the University of São Paulo (PRP-USP: 18.1.660.41.7), Brazilian Council for Scientific and Technological Development (CNPq: 302700/2016-1), Minas Gerais Research Foundation (FAPEMIG: PPM-00324-15), and Alexander von Humboldt Foundation (AvH: 3.4-8151/15037). GMFF and RBPP received scholarships from the Brazilian Coordination for the Improvement of Higher Education Personnel (CAPES) and CNPq through the Graduate School in Ecology of the Federal University of Minas Gerais (ECMVS). RLM received PhD scholarships from the São Paulo Research Foundation (FAPESP: 2015/17739-4, 2017/01816-0). SES was supported by the National Science Foundation (NSF-1456375). NL received the sandwich scholarship from CNPq and TWAS (312518/2015-3). FAR acknowledges CNPq (307974/2013-8) and FAPESP (17/50144-0 and 16/25682-5) for the financial support given for his research. FAR gratefully acknowledges support from The Leverhulme Trust for the Visiting Professorship provided.

## Author contributions

MARM conceived the project. The first version of the working question, hypothesis, and predictions was conceived by MARM together with RBPP and GMFF, and all authors contributed to improving the central argument of the study. CG and MT acquired the literature data and field data used to build the dataset of bat-plant interactions. SES reconstructed the bat phylogeny. SES and RDS built the dataset on bat morphology and performance. FAR and NL developed the new multilayer version of the centrality metrics. MARM, RLM, RBPP, GMFF, FAR, and NL performed tasks related to data analysis and coding in R and Python. The first draft of the manuscript was written by MARM, RBPP, GMFF, and RLM, and all authors contributed to editing the text.

## Competing financial interests

None to declare.

## Supplementary information

Supplementary Data 1. Dataset used to build the multilayer network, including an R code for drawing it. Available on GitHub: https://github.com/marmello77/mello-etal-2018-SD1.

Supplementary Table 1. References used to build our dataset on bat-plant interactions in the Neotropics.

Supplementary Table 2. A small dictionary of network science.

Supplementary Methods 1. Details on the calculation of centrality and the definition of the multilayer structure and the calculation of multilayer versions of the main centrality metrics.

Supplementary Methods 2. Phylogenetic and geographic signals.

Supplementary Results 1. Correlograms of centrality for each layer of the network (A: frugivory, B: nectarivory, and C: dual) and for the multilayer network (D).

Supplementary Results 2. Correlations between centrality metrics between layers. Trend lines are presented only for statistically significant relationships. (a) Correlations between three centrality metrics between layers for bat species that make interactions of frugivory and nectarivory. (b) Relationship between the centrality of bat species in the frugivory layer and the probability of being a bridge species (i.e., making dual links with the same plant species). (c) Relationship between the centrality of bat species in the nectarivory layer and the probability of being a bridge species (i.e., making dual links with the same plant species).

